# Heterogeneity in chromatin structure drives core regulatory pathways in B-cell Acute Lymphoblastic Leukemia

**DOI:** 10.1101/2024.10.04.616668

**Authors:** Arko Sen, Zhichao Xu, Sélène T. Tyndale, Jean Yasis, Chae Yun Cho, Rosalind Bump, Sahaana Chandran, Linda Luo, Yi Fu, Lillian Kay Petersen, Max Shokhirev, Dennis J. Kuo, Graham McVicker, Jesse R. Dixon

## Abstract

B-cell acute lymphoblastic leukemia (B-ALL) is the most common pediatric malignancy. Based on gene expression profiling, B-ALL can be classified into distinct transcriptional subtypes with differing disease outcomes. Many of these transcriptional subtypes are defined by mutations in transcription factors and chromatin-modifying enzymes, but how such diverse mutations lead to distinct transcriptional subtypes remains unclear. To illuminate the chromatin regulatory landscape in B-ALL, we analyzed 3D genome organization, open chromatin, and gene expression in 53 primary patient samples. At the level of 3D genome organization, we identified chromatin interactions that vary across transcriptional subtypes. These sites of variable 3D chromatin interactions correlate with local gene expression changes and are enriched for core drivers of B-ALL observed in genome-wide CRISPR knock-out screens. Sites of variable 3D genome interactions are frequently shared across multiple transcriptional subtypes and are enriched for open chromatin sites found in normal B-cell development but repressed in mature B-cells. Within an individual patient sample, the chromatin landscape can resemble progenitor chromatin states at some loci and mature B-cell chromatin at others, suggesting that the chromatin in B-ALL patient tumor cells is in a partially arrested immature state. By analyzing transcriptomic data from large cohorts of B-ALL patients, we identify gene expression programs that are shared across transcriptional subtypes, associated with B-cell developmental stages, and predictive of patient survival. In combination, these results show that the 3D genome organization of B-ALL reflects B-cell developmental stages and helps illustrate how B-cell developmental arrest interacts with transcriptional subtypes to drive B-ALL.

## Introduction

Acute lymphoblastic leukemia (ALL) is the most common pediatric cancer^1^ and, despite tremendous progress in treatment, remains the second leading cause of cancer-related death in children^2^. B-cell Acute Lymphoblastic Leukemia (B-ALL) is the most common form of ALL in children^1^. Despite a low overall mutation burden, common features of B-ALL tumors are mutations or gene fusions that affect transcriptional regulators, including transcription factors (TFs) and chromatin modifiers, as well as mutations that alter signaling pathways^3–6^. Based on gene expression, B-ALL can be grouped into about 23 transcriptional subtypes^7^, which are associated with specific mutations in transcriptional regulations (e.g., ETV6-RUNX1 fusions^8,9^, PAX5 alterations^7,10,11^, or DUX4 mutations^4,12–14^), kinase pathway alterations^15–18^ (e.g., BCR-ABL fusions^19,20^), or karyotypic abnormalities (Hyperdiploid)^19^.

The gene expression differences between transcriptional subtypes of B-ALL are potentially mediated by changes in the activity of cis-regulatory elements caused by modifications to transcription factors or chromatin modifiers. Prior studies have shown that clustering B-ALL patient samples by open chromatin mirrors the clustering into transcriptional subtypes by gene expression ^21^ and that subtype-specific open chromatin sites are associated with nearby subtype-specific differentially expressed genes^22,23^. Some subtype-specific open chromatin sites also interact with nearby gene promoters as observed by promoter capture Hi-C^22^ and some 3D genome interactions are altered during B-ALL relapse^24^. These observations suggest that changes in distal *cis* regulatory elements and 3D genome organization play a key role in B-ALL. However, despite our increasing knowledge of the chromatin landscape in B-ALL, how 3D chromatin structure varies between transcriptional subtypes in B-ALL is unknown.

Multiple transcription factor mutations associated with transcriptional subtypes affect genes that regulate normal B-cell development^25^. Mechanistically, these mutations block B-cell maturation, and in these cases, B-ALL can be thought of as a disease of altered B-cell development^26–28^. Furthermore, some transcriptional subtypes of B-ALL exhibit gene expression signatures related to B-cell development. For example, KMT2A mutated samples have gene expression patterns that resemble an early lymphocyte progenitor state^29^. Similarly, gene expression patterns in Ph+ BCR-ABL rearranged B-ALL^30^ resemble different B-cell developmental stages, suggesting a non-uniform state of developmental arrest.

To better understand chromatin organization in B-ALL and its relationship to developmental arrest, we analyzed gene expression, chromatin accessibility, and 3D chromatin architecture in primary tumor samples obtained from 53 pediatric B-ALL patients. We identified genome regions with variable 3D genome interactions across transcriptional subtypes. In contrast to what has been previously reported for gene expression and open chromatin, most of the variable 3D genome interacting regions are shared across multiple transcriptional subtypes. Genomic regions with variable chromatin interactions are enriched for essential genes in B-ALL cell lines identified in CRISPR knock-out screens. Furthermore, sites with variable 3D interactions contain open chromatin sites found in normal B-cell development but lost upon differentiation to mature B-cells. This suggests that these regions represent loci of B-lineage developmental arrest. To determine whether the signatures of developmental arrest are also found in gene expression data in patient samples, we used non-negative matrix factorization (NMF) to decompose gene expression patterns and identified 11 NMF gene expression components that are largely independent of transcriptional subtypes. Many of these components correspond to B-cell development or functional states and predict patient survival. Overall, our results shed light on the 3D chromatin state in B-ALL patient samples and reveal chromatin and gene expression programs associated with B-cell developmental arrest, which may help identify novel future therapeutic strategies.

## Results

### Chromatin profiling in B-ALL patients

To illuminate the chromatin landscape of B-ALL, we performed Hi-C, ATAC-seq, and RNA-seq on peripheral blood and bone marrow biopsies from pediatric B-ALL patients. Samples were obtained from patients at Rady Children’s Hospital, San Diego upon initial presentation of disease or relapse (Fig. 1A, Supplementary Table 1). Following stringent quality control, we obtained high-quality RNA-seq data from 53 patients, ATAC-seq data from 35 patients, and Hi-C data from 35 patients. For 13 samples, we performed deep sequencing of Hi-C libraries (346 to 804 million chromatin contacts per sample) to enable the detection of chromatin looping events at high resolution (Supplementary Table 2). Using the ATAC-seq data, we identified a median of 124,500 chromatin accessibility peaks per sample (range 104,364 to 252,875). Upon merging peaks across samples, there were 1,043,954 unique peaks highlighting the diversity in chromatin accessibility across B-ALL tumors. We also called chromatin loops in the 13 high-depth Hi-C libraries and identified a median of 21,515 loops per sample (range 8,150 to 37,190, Supplementary Table 2). After combining loops across samples and merging adjacent loop calls, 38,022 unique chromatin loops were found across these 13 samples. In addition, we identified structural variants from Hi-C data using our hic-breakfinder^31^ tool and gene fusion events from RNA-seq data using STAR-Fusion (Supplementary Tables 3 and 4).

**Figure 1:**
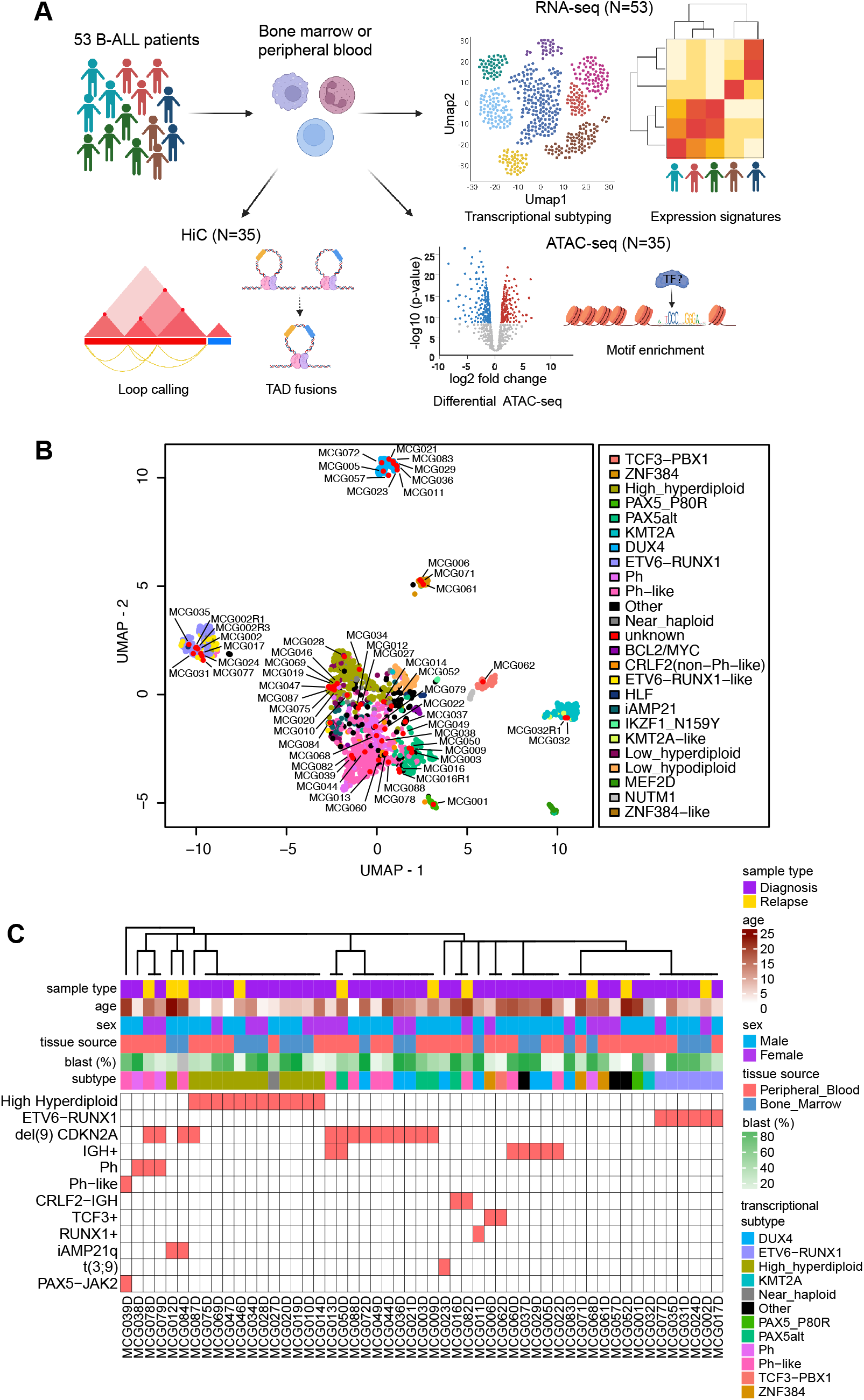
Genomic profiling of B-ALL samples. **A)** Overview of the sample collection, experiments and data analysis performed by this study. **B)** Transcriptional subtypes of B-ALL samples. Uniform manifold approximation and projection (UMAP) of gene expression data from 1,988 published RNA-seq samples colored by transcriptional subtype and 57 RNA-seq samples from this study (including matched relapse samples for a subset of the 53 total patients). B-ALL samples generated as part of this study are indicated in red and with sample labels. **C)** Clinical and demographic information of the B-ALL patient samples. **D)** RNA-seq and ATAC-seq profiles at the AGAP1 gene in eight examples samples grouped by whether they are in the DUX4 transcriptional subtype or not. **E**) Hi-C chromatin contacts surrounding AGAP1 in a DUX4 sample (MCG001) and non-DUX4 sample (MCG019).

Gene expression clustering has previously identified 23 transcriptional subtypes of B-ALL, many associated with specific mutations or gene fusions of TFs and chromatin modifiers^31^. To assign our samples to these previously identified transcriptional subtypes, we co-clustered our samples with RNA-seq profiles from 1,988 B-ALL samples (Fig. 1B). Our samples have diverse transcriptional subtypes, with 53 patient samples assigned to 12 subtypes (Fig. 1C). We compared the assigned transcriptional subtypes with the clinical karyotyping data to evaluate the consistency between classification approaches. In cases where the clinical cytogenetics identified alterations indicative of a transcriptional subtype (ETV6-RUNX1, Hyperdiploid, Ph, Ph-like), we observed a match in 24/27 cases between transcriptional subtypes and karyotype information (Supplementary Table 1). In the three cases where there was disagreement, one case (MCG012) was assigned as hyperdiploid by transcriptional clustering (defined as >51 chromosomes)^32,33^ but was labeled as 48XY by clinical karyotyping (Supplementary Table 1). A second case (MCG027) was classified as Near Haploid, but its clinical cytogenetics were ambiguous. By karyotyping, it was 52X, but chromosomal microarray assay findings were consistent with a low-hypodiploid sample. The final case (MCG016) was classified as PAX5alt from transcriptional clustering but was found to contain a CRLF2-IGH rearrangement in clinical karyotyping, which is characteristic of Ph-like samples. Interestingly, upon examining the structural variant profiles and fusion genes called from RNA-seq, we observe that this sample carries both a PAX5 fusion gene and a CRLF2-IGH rearrangement (Supplementary Figure 1F). Of the remaining 26 samples, we identified gene fusions or point mutations consistent with the classified transcriptional subtype in an additional 13 samples (Supplementary Tables 1 and 4). Of the remaining 13 samples that we could not validate the transcriptional subtype classification, three were classified as “Other” without a consistent genetic alteration, six were classified as DUX4, and four were classified as Ph-like. Notably, DUX4 mutations can be challenging to identify using short-read RNA sequencing^34^, and Ph-like samples are associated with a wide variety of kinase alterations^35^, so identifying the single causative mutation in a sample can be challenging. For both DUX4 and Ph-like, transcriptional profiling based on gene expression patterns, rather than direct detection of a somatic mutation, has previously been shown to be a robust strategy for identifying such cases^34,36,37^. These results support our classifications of patient tumor subtypes into transcriptional subtypes and highlight the value of using genomics methods to augment the cytogenetic classification of patient samples.

### Differential chromatin interactions between transcriptional subtypes are associated with core driver genes in B-ALL

To determine how 3D genome structure differs across transcriptional subtypes of B-ALL, we used the Hi-C data to identify differential chromatin interactions between subtypes at a resolution of 25kb. For this analysis, we utilized all 25kb bin pairs, not just chromatin loops, so that we could identify genome-wide differential contacts, including in regions outside of called loops. Most chromatin contacts were consistent across B-ALL subtypes, but a small subset of chromatin interactions differed between subtypes. This is similar to prior observations in T-cells, where looping is largely conserved between related cell types^38^. We identified 20,195 chromatin interactions that differ between B-ALL transcriptional subtypes, representing 0.55% (20,195/3,672,690) of tested interactions, which we call variable chromatin interactions. We compared the variable chromatin interactions with chromatin loops and found that 17% (3,404/20,195) overlapped loops in at least one patient sample.

To examine how variable chromatin interactions differ across transcriptional subtypes, we visualized them with K-means clustering (k=12, elbow method) (Fig. 2A, Supplementary Figure 2A). A subset of clusters was strongly enriched in a single transcriptional subtype or related subtypes (e.g., cluster 5 in ETV6-RUNX1, cluster 9 in Ph/Ph-like), suggesting that they may confer subtype-specific expression patterns. However, most of the variable chromatin interactions were shared between subtypes (Fig. 2A,B) and this finding was robust to the clustering method and the number of clusters (Supplemental Fig. 2B). This suggests that most variable chromatin interactions are not subtype-specific but are instead shared by multiple transcriptional subtypes.

**Figure 2:**
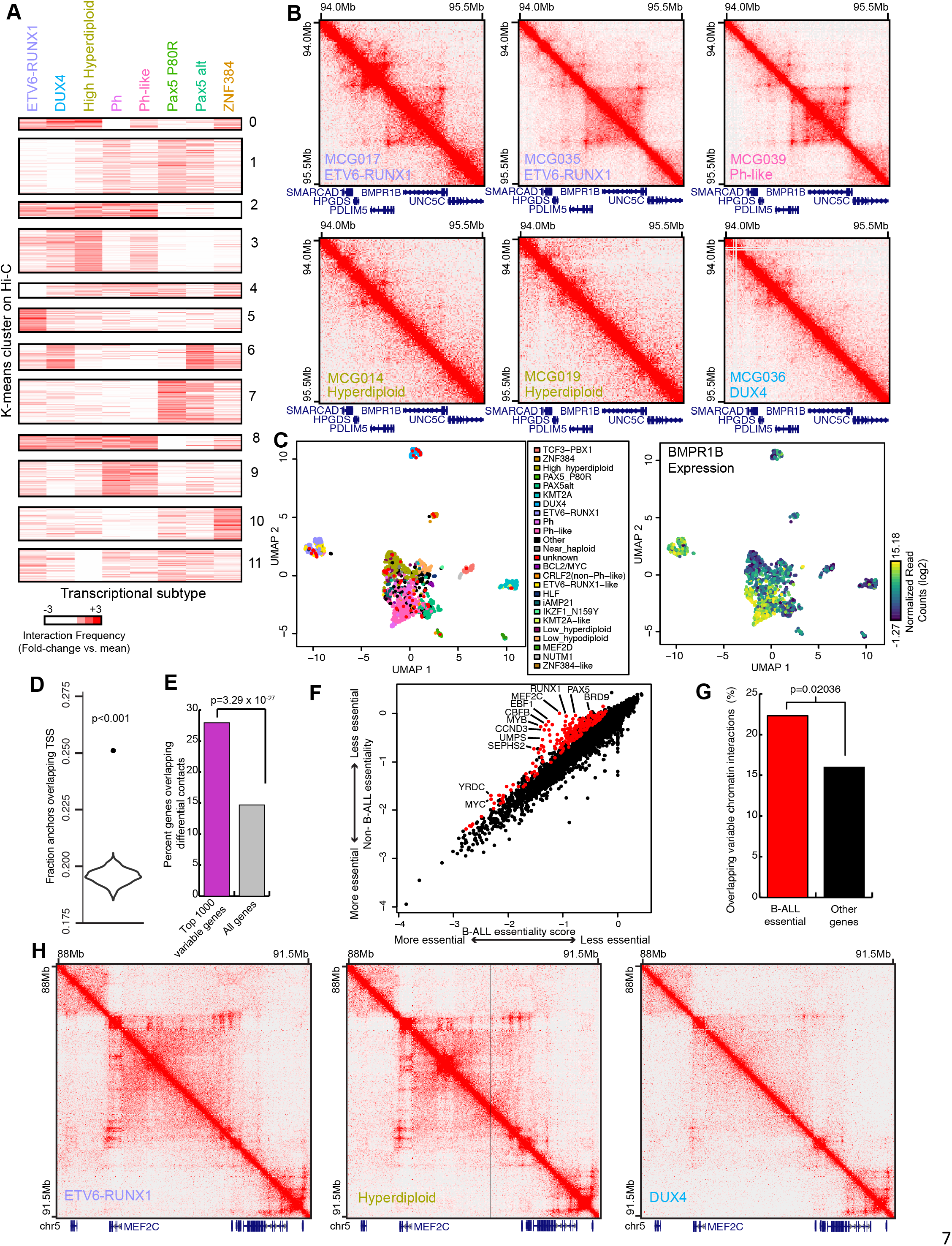
Variable chromatin contacts between transcriptional subtypes in B-ALL samples. **A)** Differential chromatin contacts between transcriptional subtypes in 25kb genomic windows. **B**) Example of a variable chromatin loop to the promoter of BMPR1B, which is present in MCG017, MCG035 and MCG039 samples from the ETV6-RUNX1 and Ph-like subtypes but absent in MCG014, MCG019 and MCG036 samples from the High Hyperdiploid and DUX4 subtypes. **C**) Uniform manifold approximation and projection (UMAP) plots colored by transcriptional subtype assignments (left panel) and the expression of BMPR1B (right panel). BMPR1B expression varies substantially and is high in samples from multiple transcriptional subtypes. **D**) Fraction of the genes with the most variable expression across subtypes that overlap differential chromatin contacts at their promoters. The distribution shows the expected overlap using randomly shuffled chromatin interactions, and the point shows the observed overlap (P < 0.001 by permutation test). **E**) Percent of genes overlapping differential 3D chromatin contacts that are classified as the top 1000 most variable genes across B-ALL samples (p=3.29×10-27, Fisher’s exact test). **F**) Average CRISPR gene essentiality scores for all genes within B-ALL cell lines (y-axis) or non-B-ALL cell lines (x-axis). A lower (more negative) essentiality score indicates a greater effect of the loss of the gene on cell fitness. Genes with significantly lower essentiality scores in B-ALL are labelled in red (FDR = 5%, Student’s T-test). **G**) Percent of B-ALL essential genes (red) or other genes (black) that overlap variable chromatin interactions (p=0.02036, Fisher’s Exact Test). **H)** Examples of Hi-C contact maps near the MEF2C gene in three patient samples. The transcriptional subtype is labelled in the lower left corner of each plot.

We examined the genomic locations of variable chromatin interactions and observed that they were frequently associated with genes. For example, we observed variable chromatin interactions in ETV6-RUNX1 and Ph-like subtypes at the *BMP1RB* gene (Fig. 2B,C), overexpression of which has been implicated in both myeloid and lymphoid leukemias^39^. Genome wide, the anchors of variable chromatin interactions were more likely to overlap with the Transcription Start Sites (TSS) of genes than expected by chance (Fig. 2D, p < 0.001, permutation test). Furthermore, the variable chromatin loops were much more likely to overlap TSSs of genes with variable expression: 27.9% of the top 1000 most variable genes overlapped with variable chromatin interaction anchors, compared with 14.7% of all genes (P = 3.26 × 10^−27^, Fisher’s exact test) (Fig. 2E). Thus, variable chromatin loops are associated with variable gene expression.

Variability in chromatin structure and gene expression may contribute to phenotypic differences between transcriptional subtypes or may instead reflect biological “noise” that is irrelevant to B-ALL pathogenesis. To determine whether variable chromatin interactions were associated with core regulatory pathways in B-ALL, we analyzed essential genes in B-ALL using data from CRISPR/Cas9 knock-out screens. Specifically, we computed the average dependency scores for genes using B-ALL cell lines and non-B-ALL cell lines using data from the DepMap CRISPR/Cas9 essentiality screens project^40^ (Fig 2F). We identified a set of “core B-ALL driver genes” with significantly different essentiality scores between B-ALL and non-B-ALL cell lines, with more negative scores indicating greater effects on cell survival in B-ALL. In total, 202 genes had significantly greater effects on cell survival in B-ALL cell lines compared with non-B-ALL cell lines (t-test, FDR 5%, difference in essentiality scores <-0.2), including several known regulators of B-ALL and B-cell development like *PAX5, EBF1*, and *MYC*. To determine whether the variable chromatin interaction regions are associated with the core driver genes of B-ALL, we compared the B-ALL core driver genes with variable chromatin interaction anchors. We observed that 21.8% of B-ALL core driver genes were associated with variable chromatin interactions compared to 16.0% of non-B-ALL essential genes (Fig. 2G) (p=0.02, Fisher’s Exact Test). One example is *MEF2C*, which is one of the most selectively essential B-ALL genes and which has highly variable chromatin interactions across patient samples (Fig. 2H). Taken together, these results suggest that variable chromatin interactions regulate core genes that are essential for B-ALL cell survival.

### Variable chromatin interactions are associated with a B-lineage partial developmental arrest

Given the association between variable chromatin interactions and core driver genes in B-ALL, we were interested in understanding the molecular basis for variability in chromatin interactions across transcriptional subtypes. Examining both the Hi-C and ATAC-seq data, we observed that variable chromatin interactions were often associated with changes in open chromatin (Fig. 3A). To quantify this, we computed the correlation between Hi-C contact frequency and normalized ATAC-seq signal over all ATAC-seq peaks within the anchors of variable chromatin interactions (Fig. 3B). At most variable interaction loci, the correlations are positive and 1,427 unique ATAC-seq peaks were significantly correlated with at least one variable 3D chromatin interaction (empirical FDR = 5%, n=1000 permutations).

**Figure 3:**
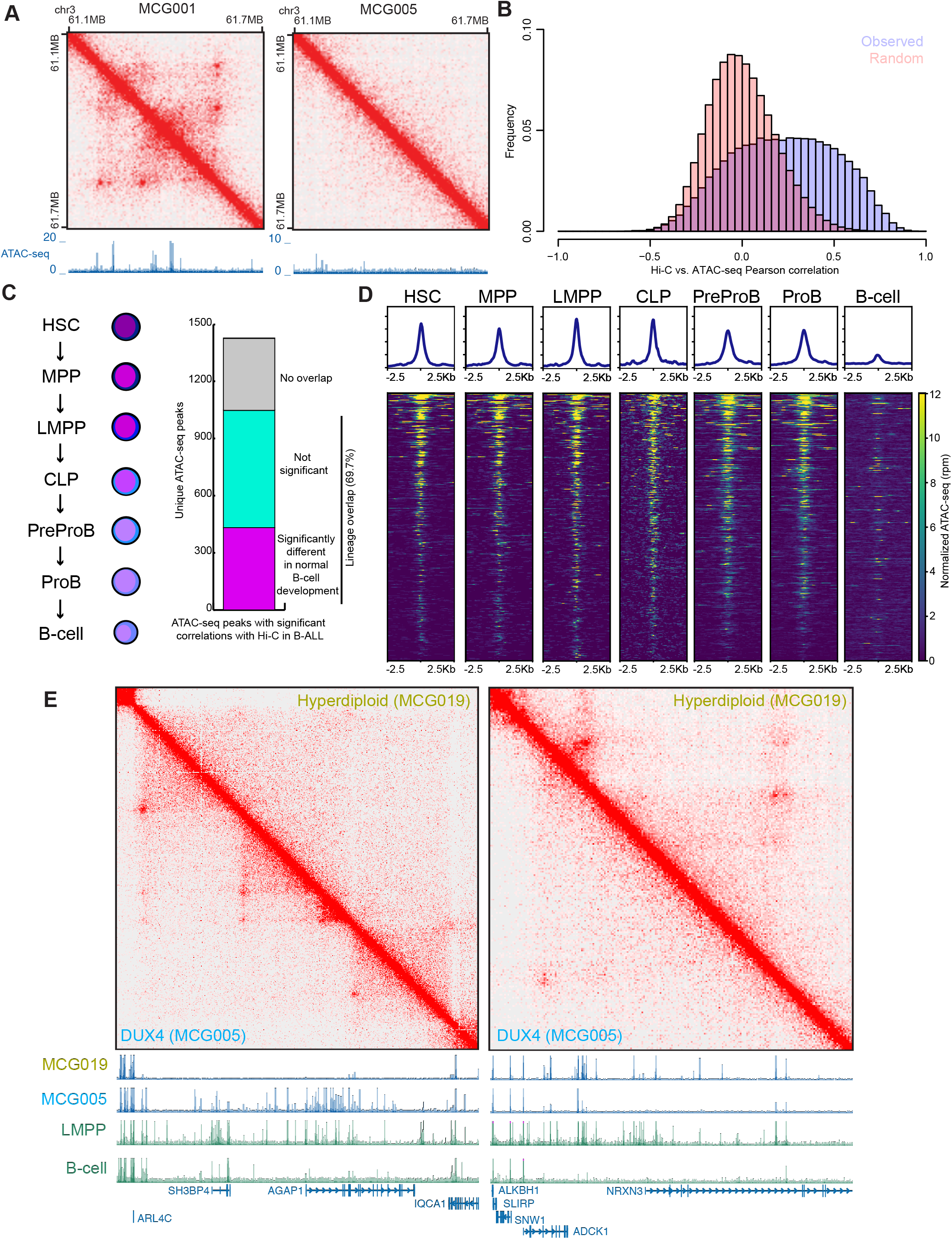
Variable chromatin interactions are associated with open chromatin from B-lineage development. **A)** Example of differential 3D genome interactions between two samples (MCG001 - PAX5_P80R and MCG005 DUX4). Shown below is ATAC-seq data from the same patients at the same locus. **B**) Histograms of Pearson correlation coefficients between chromatin interaction frequency and open chromatin signal (ATAC-seq) at sites of variable chromatin interactions. The purple histogram shows the true observed Pearson correlations while the pink histogram shows the Pearson correlations when the vector of ATAC-seq data is randomly permutated. The permutation is performed 1000 times. **C**) Diagram showing B-lineage developmental cells with ATAC-seq data (left) and the fraction of ATAC-seq peaks with significant correlations with Hi-C data that overlap these normal B-lineage developmental ATAC-seq data. Overlapping peaks are distinguished as being significantly different among normal B-lineage cells (pink) or not significantly different (green) (Quasi-likelihood F-test, edgeR, FDR = 5%). **D)** ATAC-seq open chromatin profiles for B-ALL 3D genome correlated ATAC-seq peaks in normal B-lineage cell development. **E)** Examples of differential 3D genome interactions in two patients samples (MCG005 - DUX; MCG019 - Hyperdiploid) and the associated ATAC-seq data at the same locus. ATAC-seq data is shown for each patient (top two tracks) as well as for cells in normal B-lineage development (LMPP - Lineage Primed Multi-potent Progenitor).

To determine whether the 1,427 ATAC-seq peaks that correlate with 3D genome structure are cancer-specific or found in the normal development of B-cells, we downloaded publicly available B-cell lineage development ATAC-seq data spanning hematopoietic stem cells (HSC) to mature B-cells^41,42^ (Fig. 3C). We identified a consensus list of 195,729 ATAC-seq peaks from normal B-cell lineage development and compared them with the 1,427 ATAC-seq peaks that correlated with 3D genome interactions in B-ALL. We found that 69.7% (n=995/1,427) of the B-ALL ATAC-seq peaks are found in normal B-cell development (Fig. 3C). Furthermore, 38.0% (378/995) of these peaks were significantly different between developmental lineages (FDR = 1% quasi-likelihood F-test, 4-fold difference from mean). This suggests that sites with variable open chromatin and 3D genome interactions are enriched at developmentally regulated B-cell loci.

To determine whether the variable B-ALL open chromatin sites reflect specific developmental stages, we compared the ATAC-seq signal from normal B-cell development at the 1,427 B-ALL ATAC-seq peaks that were correlated with 3D genome structure (Fig. 3D). While no single developmental stage was strongly enriched above others for open chromatin signal, there was a reduction in ATAC-seq signal in mature B-cells (Fig. 3D). Thus the variable ATAC-seq peaks in B-ALL are present in B-cell progenitors but lost in mature B-cells, consistent with B-ALL resulting from an arrest of normal B-cell development. Furthermore, the fact that variable chromatin interactions are found in different transcriptional subtypes in different clusters (Fig. 2A) suggests that the B-ALL samples do not uniformly represent the progenitor B-cell stages. Instead, a given B-ALL sample may be more similar to B-cell progenitors at some loci but be more similar to mature B-cells at others. For example, when we compare ATAC-seq signals for a DUX4 sample (MCG005) and a Hyperdiploid sample (MCG019) at the *AGAP1* locus, the DUX4 sample more closely resembles progenitor cells, and the Hyperdiploid sample resembles mature B-cells (Fig. 3E, left). However, at the *NRXN3* locus, the opposite is true—here, the DUX4 sample resembles mature B-cells, while the Hyperdiploid sample resembles progenitor cells (Fig. 3E, right). These results suggest that B-ALL cells may not reflect a complete B-cell developmental arrest. Instead, they may have partial developmental arrest that, at the chromatin level, is locus specific and related to the transcriptional subtype.

### Shared regulatory programs across transcriptional subtypes

Given our observations that the variable open chromatin and 3D genome interactions reflect chromatin states of normal B-cell progenitors, we wanted to test whether this phenomenon could be observed more broadly in the gene expression patterns of B-ALL patient samples. Since the similarity with progenitor cell ATAC-seq signals differs across loci, we reasoned that combinations of multiple gene expression programs would better describe each sample’s gene expression and chromatin patterns. We therefore performed non-negative matrix factorization (NMF)^43^ to decompose the gene expression of each sample into multiple transcriptional programs. NMF is well suited to this task as it can identify essential components of high-dimensional data while allowing each sample to be comprised of more than one component. NMF has previously been used to decompose tumor mutation patterns into mutational signatures^44^, to discover molecular subgroups for tumors such as medulloblastoma^45,46^, and to define regulatory programs from chromatin accessibility^47^ and single-cell gene expression^48^. Recently, NMF was employed to detect B-cell developmental programs within the Ph-like B-ALL transcriptional subtype^45^.

When applied to gene expression, NMF factorizes the gene expression matrix into two non-negative matrices: a basis matrix that describes the contribution of each gene to each NMF component and a sample matrix that describes the contribution of each component to each sample (Fig. 4A). We applied NMF to gene expression data from 1,266 B-ALL transcriptomes^46^, decomposing each sample into its constituent NMF components^47^. To specify the factorization rank (k) and the number of components, we tested values between k=6 and k=24. We chose k=11 because this resulted in a cophenetic coefficient ≥ 90%, indicating stable clustering^48^, and low pairwise correlations between components (<20%), indicating non-redundancy(Suppl Fig. 3A, B). We then used the basis matrix from the larger panel to assign NMF components and derive a sample matrix for the smaller panel using QR factorization (Fig. 4B).

**Figure 4:**
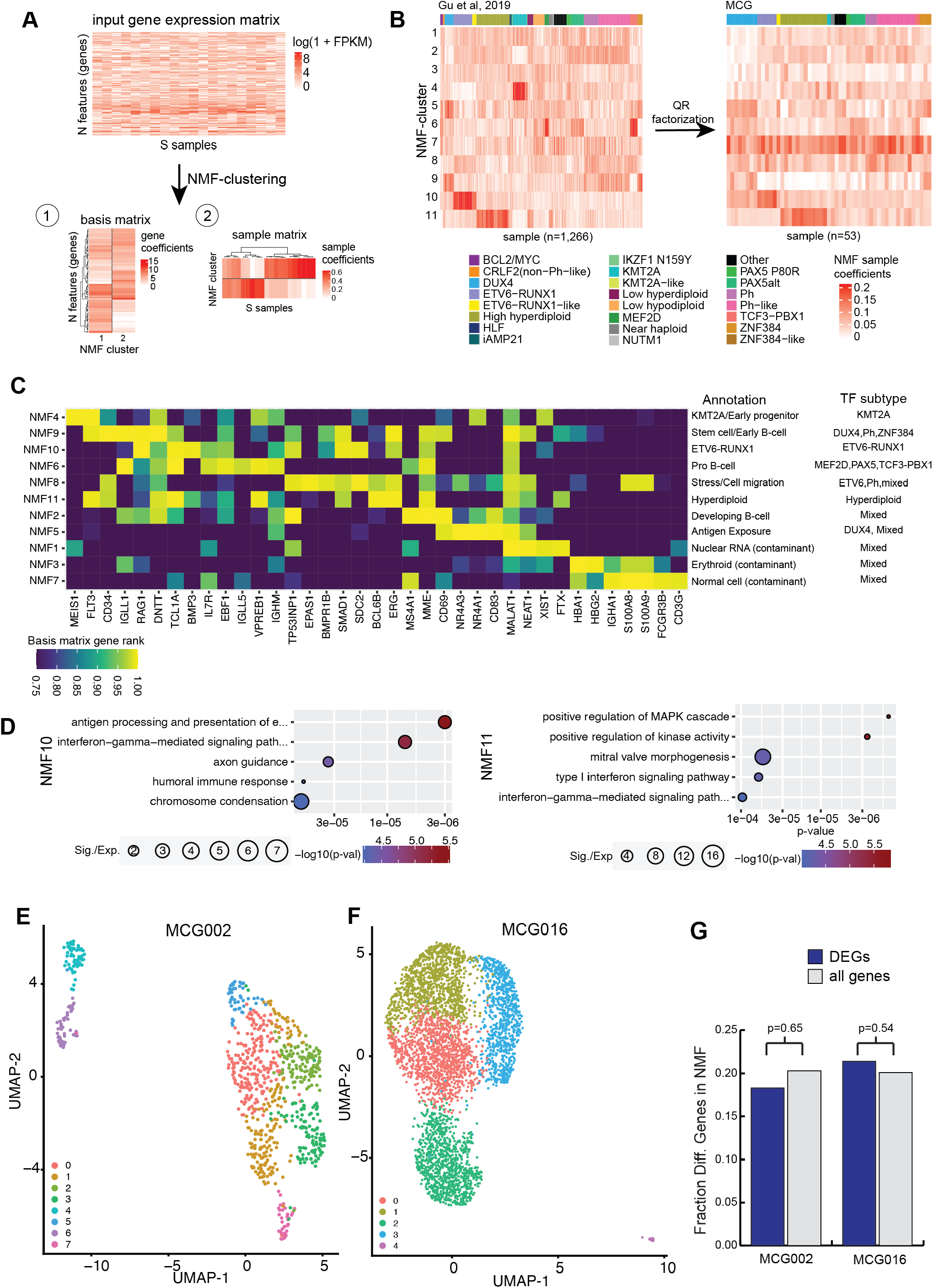
Latent regulatory programs revealed with NMF. **A)** Example diagram for non-negative matrix factorization (NMF) to decompose a gene expression matrix into K gene expression programs, described by a basis matrix and a sample coefficient matrix. The basis matrix defines the contribution of each gene’s expression to each gene expression program. The sample matrix defines the contribution of each gene expression program to each sample’s overall gene expression profile. **B**) Sample coefficient matrix for a reference data set of 1,266 B-ALL samples with published gene expression data (left panel). The samples are labeled and grouped by their previously established transcriptional subtypes. The sample coefficient matrix for 53 B-ALL patients from this study (right panel), which we derived using QR factorization. **C)** Heatmap showing the rank of each gene in the basis matrix for different NMF components for select marker genes of hematopoiesis. **D**) Gene Ontology analysis for genes in the top 100 of the basis matrices for NMF10 and NMF11. **E**) UMAP plot of single-cell RNA-seq data from the MCG002 patient. Points are labeled by cluster. **F**) UMAP of single-cell ATAC+RNA-seq multi-ome from the MCG016 patient. Points are labeled by cluster. **G**) The fraction of differentially expressed genes between B-ALL clusters in the MCG002 and MCG016 samples that overlap genes in the top 100 of the basis matrix of any NMF signature. P-values are from the Fisher’s exact test.

To illuminate the transcription programs identified by NMF, we examined the top-scoring genes in the basis matrix for each component and the association of each component with different transcriptional subtypes (Fig. 4B, C). Some of the NMF components correspond to a single transcriptional subtype. For example, components 4, 10, and 11 correspond to the KMT2A, ETV6-RUNX1, and Hyperdiploid transcriptional subtypes. The top-scoring genes in the basis matrix for these components also suggest distinct functions or developmental stages. For example, the early hematopoietic developmental genes *CD34* and *MEIS1* are two of the top-scoring genes for the NMF4 (KMT2A associated) component, suggesting that it corresponds to an early lymphoid progenitor state, which has been recently proposed in KMT2A B-ALL^29^. Gene Ontology analysis of the top 500 genes in the NMF10 (ETV6-RUNX1 associated) component revealed a strong enrichment for terms related to interferon signaling and antigen presentation. Similarly, the top 500 genes for the NMF11 (Hyperdiploid associated) were enriched for MAP kinase signaling (Fig. 4D, Supplementary Table 5).

However, Other NMF components represent samples across multiple subtypes. Three of these components, NMF1, NMF3, and NMF7, appear to reflect technical artifacts of experimentation and sample collection. The top genes for NMF3 and NMF7 are markers of non-tumor cells: NMF3 is strongly enriched for erythrocyte-specific genes, and NMF7 is enriched for neutrophil, T-cell, and mature B-cell genes (Fig. 4C, Supplementary Table 5). Furthermore, unlike other components, NMF7 has a significant negative correlation with the percentage of blasts in the sample at the time of collection (Supplementary Fig. 3C, Pearson r= -0.76, p=6.29×10^−6^). Similarly, we suspect that NMF1 is related to sample integrity. GO term analysis shows enrichment for multiple terms related to RNA processing and nuclear bodies, such as Nuclear Speckles, Cajal bodies, and U4/U6 x U5 snRNP complex (Supplementary Table 5), and many of the top genes for this component are nuclear-retained RNAs, such as *MALAT1, NEAT1, XIST*, and *FTX*. The association with *XIST* and *FTX* initially suggested that NMF1 might be related to sex. However, NMF1 is not enriched in female samples. Instead, our interpretation is that NMF1 is related to technical aspects of sample processing because nuclear RNAs are likely to have a higher relative abundance in samples with more lysis or fragmentation.

The remaining NMF components (NMF2, NMF5, NMF6, NMF8, and NMF9) are not explicitly enriched in a single transcriptional subtype and do not appear to reflect technical artifacts. Instead, these components are enriched for genes that are expressed at different stages of B-cell development. NMF9 reflects early B-cell development, with the pre-pro B cell and pro-B cell genes^49^ *CD34, IGLL1, RAG1*, and *DNTT* (encoding TdT) all within the top 10 highest-ranked genes in the basis matrix (Fig. 4C). Continuing through B-cell development, NMF6 displays signatures of developing B-cells, with relative enrichment of pro-B cell and pre-B-cell genes such as *VPREB1, EBF1*, and *IL7R* (Fig. 4C). Furthermore, NMF6 shows a strong enrichment for cycling cells by GO term analysis (Supplementary Table 5) and is the only NMF component with the cycling marker *MKI67* in the top 100 genes. In contrast, NMF2 shows signatures of later B-cell development with enrichment for the late pre-B cell or immature B-cell marker *MS4A1* (encoding CD20). NMF5 shows signatures of antigen exposure, as its top two genes are the orphan nuclear receptors *NR4A1* and *NR4A3*, which are rapidly induced in response to antigen^50^. NMF5 also shows enrichment for *CD83*, a marker for activated B-cells^51^. NMF8 expresses some early B-cell development genes, such as *CD34*, but also expresses markers of stress and cell migration, including *S100A8* and *S100A9*^52^. In summary, normal B cell development signatures are common in primary patient B-ALL samples and across transcriptional subtypes.

Differences in the NMF components associated with B-ALL patient samples could reflect intra-tumoral heterogeneity rather than consistent changes in gene expression programs shared by all tumor cells within a sample. To address this, we performed single-cell RNA sequencing on one patient sample (MCG002) and multi-omic single-cell RNA-seq and single-cell ATAC-seq assays on another patient sample (MCG016) (Fig. 4E,F). For both samples, we performed clustering, embedding in low dimensional space, and called differentially expressed genes using Seurat. By performing marker gene analysis, we identified clusters that were of non-B-cell origin (clusters 4 and 6 for MCG002, cluster 4 for MCG016) and one cluster in each sample that was defined by cycling cells based on the expression of marker gene *MKI67* (Supplementary Figure 4). The remaining clusters in both samples were of B-cell origin and had high expression of B-cell marker genes, such as *EBF1* (Supplementary Figure 4). These B-cell clusters likely represent tumor blasts, as for both samples, the blast percentages obtained when the samples were isolated were high (55% for MCG002, 96% for MCG016).

To determine whether NMF components reflect intra-tumoral heterogeneity, we identified genes that varied across the non-cycling B-cell clusters by identifying marker genes that were upregulated 2-fold relative to the remaining clusters (FDR 5%, Wilcoxon Rank Sum test). We identified 126 genes that differed across B-cell clusters in MCG002 and 299 that differed across B-cell clusters of MCG016. We then compared the overlap of these genes with the top 100 basis-matrix genes for any non-artifactual NMF component (i.e. excluding NMF1, 3, 7). For both MCG002 and MCG016, the differentially expressed marker genes were not more likely to overlap the top NMF component genes compared to non-differentially expressed genes (Fig. 4G, p=0.65 for MCG002 and p=0.54 for MCG016, Fisher’s exact test). This argues that NMF components are not explained by intra-tumoral heterogeneity and are instead more likely to be due to cell-intrinsic expression signatures that mirror aspects of B-cell developmental or functional states.

### NMF components are associated with variable chromatin interactions and predict patient survival

As we noted above, variable chromatin interactions between B-ALL patients were frequently shared across different transcriptional subtypes (Fig. 2C, D), prompting us to investigate shared gene expression components in B-ALL using NMF. We next wanted to test more directly whether the NMF components contribute to the shared patterns of chromatin interactions.

First, we identified genes with expression that correlated with variable chromatin interactions (Fig. 2A) overlapping their promoters. We computed Pearson correlation between gene expression and chromatin interaction frequency and identified 780 genes with expression that was significantly correlated with chromatin interaction frequency at the promoter (Fig. 5A, B) (FDR = 5%, by permutation test; R>0.318). We then asked if these genes overlapped with the top 100 genes in the basis matrix for the NMF components. NMF6, 8, 10, and 11 showed significant enrichments and, in each of these cases, over 20% of the top NMF genes had significant correlations between expression and promoter chromatin interactions (Fig. 5C). While NMF10 and 11 specifically correspond to the ETV6-RUNX1 and Hyperdiploid subtypes, NMF6 and 8 are shared across multiple transcriptional subtypes (Fig. 3B, F), suggesting that these expression programs may underly the variable chromatin interactions that are shared across distinct transcriptional subtypes.

**Figure 5:**
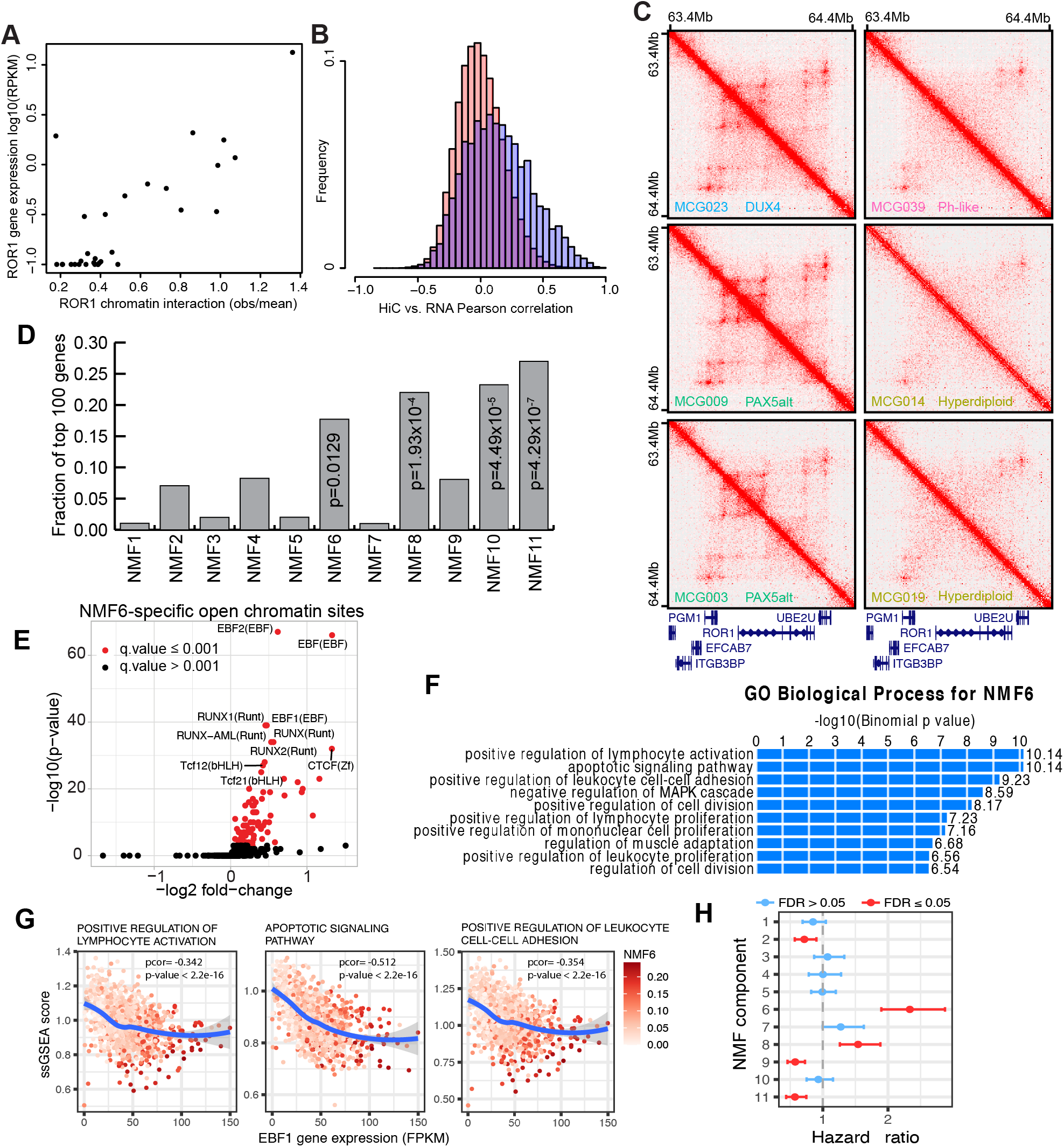
NMF components correlate with 3D genome organization and predict patient survival. **A)** Example scatter plot showing the correlation of 3D chromatin interactions (x-axis) and RNA-seq data (y-axis) for contacts at the ROR1 gene transcription start site. ROR1 is in the top 100 genes in the basis matrix for NMF signature 6. **B**) Histograms of Pearson correlation coefficients between chromatin interaction frequency and RNA-seq at sites of variable chromatin interactions. The blue histogram shows the true observed Pearson correlations while the pink histogram shows the Pearson correlations when the vector of RNA-seq data is randomly permutated. The permutation is performed 1000 times. **C**) 3D genome contact maps in six patients at the ROR1 gene locus showing variable 3D genome interactions. **D)** Fraction of genes in the top 100 basis matrix for each NMF gene signature that shows significant correlations with 3D genome interactions at their transcription start sites. NMF components 6, 9, 10, and 11 all show significant correlations (Fisher’s exact test). **E**) Motif enrichment of known TF motifs at NMF6-specific open chromatin sites. **F**) The Genomic Regions Enrichment of Annotations Tool (GREAT) analysis of ATAC-seq bins associated with NMF6. **G)** Scatter plot for correlation between single sample GSEA scores for top 3 pathways from Fig. 5F and EBF1 FPKM-normalized gene expression for the Gu et al. dataset. **H)** Hazard ratio (i.e., Cox regression model coefficients) for normalized NMF component sample values. Points and error bars are colored based on their FDR-corrected p-value from Cox regression survival analysis.

To identify the cis-regulatory elements that may underlie NMF components, we divided ATAC-seq peaks into uniform 1Kb bins and correlated chromatin accessibility in the ATAC-seq bins with the NMF coefficients for each sample. We identified 31,539 significantly correlated bins under an FDR of 1% (Supplementary Figure 5A, B, Supplementary Table 6). To determine which TFs may be driving the NMF components, we compared the frequency of TF motifs in the positively correlated bins for each NMF component to their frequency in 10,000 randomly selected ATAC-seq bins. Under a stringent FDR of 0.001, we identified 202 known vertebrate motifs from the JASPAR database that were associated with at least one NMF component. Many of the NMF components were enriched for hematopoietic TF motifs, such as those in the ETS, EBF, and STAT families (Supplementary Figure 5C). Given their association with variable 3D chromatin interactions, we focused our further analysis on NMF6 and 8. NMF8 is broadly expressed across many transcriptional subtypes and is enriched for ETS motifs (Supplementary Figure 5D and Supplementary Table 7). NMF6 is predominantly expressed in the BCL2/MYC, MEF2D, Pax5Alt, and TCF3:PBX1 transcriptional subtypes. It is enriched for several motifs in the RUNX, Tcf, CTCF, and early B cell factor (EBF) families (Fig 5D and Supplementary Table 7).

The enrichment of EBF motifs in NMF6 is interesting because *EBF1* is critical for early B-cell development^53–55^, mutations in *EBF1* are frequent in B-ALL^10^, and NMF6 is enriched for genes related to early B-cell development, including *EBF1* (Fig. 4C). GREAT analysis of 1,819 open chromatin bins positively correlated with NMF6 coefficients showed enrichment for B-cell developmental processes, such as lymphocyte activation, apoptotic signaling, and leukocyte cell-cell adhesion pathways (Fig. 5E). Moreover, we observed a negative correlation between *EBF1* expression and single-sample gene set enrichment (ssGSEA) scores for these pathways (Fig. 5F). The reduced apoptotic signaling and lymphocyte activation are characteristic of rapidly proliferating cells and often indicate a less differentiated cell state. These results are consistent with the notion that EBF1 is important for maintaining the repression of B-cell maturation and differentiation genes^56–59^. These results also imply that the hidden transcriptional signatures we identified through NMF may be driven, at least in part, by varying patterns of expression of TFs that are crucial regulators of B-cell development.

Given that sites of variable chromatin interactions are enriched for B-cell developmental gene signatures (Fig. 5C) and core driver genes in B-ALL (Fig. 2F), we wanted to determine whether the developmental gene expression signatures identified using NMF are predictive of patient survival. To this end, we analyzed 204 pediatric B-ALL samples that have both gene expression and survival data in the Therapeutically Applicable Research to Generate Effective Treatments (TARGET, phase II) database^60^. Using the same QR factorization approach that we used to assign NMF components to our samples (Fig. 4B), we assigned NMF components to the TARGET samples. For each NMF component, we then performed a multivariate Cox regression of event-free survival against the standardized NMF sample values. As covariates, we included standardized values for age, white blood cell count at diagnosis, sex, and central nervous system (CNS) status. Three NMF components, NMF2, 9, and 11, had significant negative hazard ratio coefficients, indicating they are associated with increased survival (Fig. 5G). NMF component 11 is enriched for the Hyperdiploid transcriptional subtype (Fig. 3B, F), which is consistent with Hyperdiploidy being associated with lower risk B-ALL^61,62^. NMF2 and NMF9 express mature (NMF2) and early (NMF9) B-cell developmental genes, such as MS4A1/CD20 (NMF2) and FLT3/CD34 (NMF9). In contrast, two components, NMF6 and NMF8, had significant positive hazard ratios, indicating worse survival (Fig. 5G). Thus, NMF components are predictive of patient survival in B-ALL.

## Discussion

Transcriptional subtypes of B-ALL are associated with distinct prognostic outcomes and treatment regimens^61^. Therefore, a better understanding of the differences in their chromatin architecture and how it regulates gene expression can inform putative gene targets for therapeutic interventions. In this study, we analyzed gene expression, open chromatin, and 3D genome organization across 53 primary patient B-ALL samples. We identified gene regulatory programs (NMF components) that are shared across transcriptional subtypes and that contribute to variability in 3D genome organization. These shared regulatory programs reflect differences in B-cell developmental stages, are predictive of patient survival, and represent a previously underappreciated aspect of variability in B-ALL pathophysiology.

Somatic mutations that define B-ALL are highly enriched within transcription factors and chromatin-modifying enzymes^3,10^. Therefore, it is logical that B-ALL samples cluster into diverse transcriptional subtypes associated with these mutations based on gene expression measured by RNA-seq. In contrast, however, our data show that at the level of 3D genome organization, only a minority of the chromatin interactions that vary across transcriptional subtypes are specific to a given subtype. This suggests that the mutant TFs and chromatin modifiers that define B-ALL mostly do not reprogram 3D genome organization alone. Instead, these factors act on shared 3D genome structural features that are associated with differences in normal B-cell physiology.

Sites with variable chromatin interactions are enriched for core drivers of B-ALL physiology from CRISPR screens^40^, suggesting that different tumors have different critical dependencies. In addition, variable chromatin interactions are enriched at open chromatin sites that are specific to B-cell progenitor lineages but largely repressed in mature B-cells. This suggests that these genome regions reflect a block in differentiation in B-lineage development, a phenomenon previously suggested to play a vital role in B-ALL^25,28^. Furthermore, the regions of the genome that retain the progenitor chromatin states vary between patient samples. This raises an exciting possibility for future treatment strategies: specific pathways could be targeted that selectively retrain progenitor chromatin states at specific loci and push B-ALL cells into a differentiated state. This will require future studies to better characterize the factors that drive locus-specific developmental programs.

Our study also highlights the power of genomic methods for classifying and stratifying patient tumor samples. Genomics methods such as RNA-seq are powerful tools for identifying somatic mutations like gene fusions and can also be used to infer the transcriptional states in individual samples based on larger reference panels. For example, we can classify our primary patient tumor samples into known transcriptional states using public reference panels. Further, our NMF analysis can identify novel expression signatures that predict patient survival. Specifically, we have identified signatures of NMF6 and NMF8 as associated with adverse prognostic outcomes. Understanding the molecular features that define the worse prognosis in these patients will yield important insights into B-ALL physiology and help identify new personalized treatment strategies.

## Acknowledgements

We would like to thank and acknowledge the patients and families at Rady Children’s Hospital San Diego for participation in this study. We would also like to acknowledge the Rady Children’s Institute for Genomic Medicine Biorepository. This study was initiated using a grant from the Padres Pedal the Cause foundation (PTC2017; Curebound) to J.R.D., G.M., and D.J.K. This work was supported by a grant to J.R.D. from the 4D Nucleome project (U01CA260700). This work was supported by the National Cancer Institute funded Salk Institute Cancer Center (P30-CA014195) supporting the Salk Next Generation Sequencing Core and the Salk Bioinformatics Core. We would like to acknowledge Nasun Hah, Manching Ku, and Ling Ouyang from the Salk Next Generation Sequencing core for help with sample sequencing. We would like to acknowledge April Williams and Max Shokhirev from the Salk Bioinformatics for with help with initial data processing.

## Author contributions

J.R.D., G.M., and D.J.K. conceived the study. A.S. was the lead data analyst. Z.X. led the experimental efforts and contributed to data analysis. D.J.K. led the clinical sample collection and patient recruitment. L.L., S.T.T., J.Y., C.Y.C, R.B., S.C. led the sample isolation and purification. Z.X., S.T.T., J.Y., C.Y.C., R.B., S.C. performed experimental assays and generated sequencing libraries. A.S., Z.X., R.B., Y.F., L.K.P., and M.S. contributed to data analysis. A.S., Z.X., J.R.D., and G.M. wrote the manuscript.

## Competing Interests

The authors declare no competing interests.

## Data availability

The data generated in this study is available in dbGaP accession phs003226.

## Notes

### Competing Interest Statement

The authors have declared no competing interest.

## References

1. Siegel, R. L., Giaquinto, A. N. & Jemal, A. Cancer statistics, 2024. CA Cancer J. Clin. 74, 12–49 (2024).

2. Curtin, S. C. & Anderson, R. N. Declines in Cancer Death Rates among Children and Adolescents Less than 20 Years of Age in the United States, 2001 to 2021. http://dx.doi.org/10.15620/cdc:134499 (2023) doi:10.15620/cdc:134499.

3. Brady, S. W. et al. The genomic landscape of pediatric acute lymphoblastic leukemia. Nat. Genet. 54, 1376–1389 (2022).

4. Liu, Y. F. et al. Genomic Profiling of Adult and Pediatric B-cell Acute Lymphoblastic Leukemia. EBioMedicine 8, 173–183 (2016).

5. Iacobucci, I. & Mullighan, C. G. Genetic basis of acute lymphoblastic leukemia. J. Clin. Oncol. 35, 975–983 (2017).

6. Teitell, M. A. & Pandolfi, P. P. Molecular genetics of acute lymphoblastic leukemia. Annu. Rev. Pathol. 4, 175–198 (2009).

7. Gu, Z. et al. PAX5-driven subtypes of B-progenitor acute lymphoblastic leukemia. Nat. Genet. doi:10.1038/s41588-018-0315-5.

8. Romana, S. P. et al. The t(12;21) of acute lymphoblastic leukemia results in a tel-AML1 gene fusion. Blood 85, 3662–3670 (1995).

9. Shurtleff, S. A. et al. TEL/AML1 fusion resulting from a cryptic t(12;21) is the most common genetic lesion in pediatric ALL and defines a subgroup of patients with an excellent prognosis. Leukemia 9, 1985–1989 (1995).

10. Mullighan, C. G. et al. Genome-wide analysis of genetic alterations in acute lymphoblastic leukaemia. Nature 446, 758–764 (2007).

11. Kuiper, R. P. et al. High-resolution genomic profiling of childhood ALL reveals novel recurrent genetic lesions affecting pathways involved in lymphocyte differentiation and cell cycle progression. Leukemia 21, 1258–1266 (2007).

12. Yasuda, T. et al. Recurrent DUX4 fusions in B cell acute lymphoblastic leukemia of adolescents and young adults. Nat. Genet. 48, 569–574 (2016).

13. Lilljebjörn, H. et al. Identification of ETV6-RUNX1-like and DUX4-rearranged subtypes in paediatric B-cell precursor acute lymphoblastic leukaemia. Nat. Commun. 7, (2016).

14. Zhang, J. et al. Deregulation of DUX4 and ERG in acute lymphoblastic leukemia. Nat. Genet. 48, 1481–1489 (2016).

15. Den Boer, M. L. et al. A subtype of childhood acute lymphoblastic leukaemia with poor treatment outcome: a genome-wide classification study. Lancet Oncol. 10, 125–134 (2009).

16. Roberts, K. G. et al. Genetic alterations activating kinase and cytokine receptor signaling in high-risk acute lymphoblastic leukemia. Cancer Cell 22, 153–166 (2012).

17. Mullighan, C. G. et al. Deletion ofIKZF1and prognosis in acute lymphoblastic leukemia. N. Engl. J. Med. 360, 470–480 (2009).

18. Tasian, S. K., Loh, M. L. & Hunger, S. P. Philadelphia chromosome–like acute lymphoblastic leukemia. Blood 130, 2064–2072 (2017).

19. Ribeiro, R. C. et al. Clinical and biologic hallmarks of the Philadelphia chromosome in childhood acute lymphoblastic leukemia. Blood 70, 948–953 (1987).

20. Rowley, J. D. A new consistent chromosomal abnormality in chronic myelogenous leukaemia identified by quinacrine fluorescence and giemsa staining. Nature 243, 290–293 (1973).

21. Diedrich, J. D. et al. Profiling chromatin accessibility in pediatric acute lymphoblastic leukemia identifies subtype-specific chromatin landscapes and gene regulatory networks. Leukemia (2021) doi:10.1038/s41375-021-01209-1.

22. Barnett, K. R. et al. Epigenomic mapping reveals distinct B cell acute lymphoblastic leukemia chromatin architectures and regulators. Cell Genom. 3, 100442 (2023).

23. Wang, H. et al. Chromatin accessibility landscape of relapsed pediatric B-lineage acute lymphoblastic leukemia. Nat. Commun. 14, (2023).

24. Narang, S. et al. Clonal evolution of the 3D chromatin landscape in patients with relapsed pediatric B-cell acute lymphoblastic leukemia. Nat. Commun. 15, 7425 (2024).

25. Somasundaram, R., Prasad, M. A. J., Ungerbäck, J. & Sigvardsson, M. Transcription factor networks in B-cell differentiation link development to acute lymphoid leukemia. Blood 126, 144–152 (2015).

26. Jia, Z. & Gu, Z. PAX5 alterations in B-cell acute lymphoblastic leukemia. Front. Oncol. 12, (2022).

27. Liu, G. J. et al. Pax5 loss imposes a reversible differentiation block in B-progenitor acute lymphoblastic leukemia. Genes Dev. 28, 1337–1350 (2014).

28. Fischer, U. et al. Cell fate decisions: The role of transcription factors in early B-cell development and leukemia. Blood Cancer Discov. 1, 224–233 (2020).

29. Khabirova, E. et al. Single-cell transcriptomics reveals a distinct developmental state of KMT2A-rearranged infant B-cell acute lymphoblastic leukemia. Nat. Med. 28, 743–751 (2022).

30. Kim, J. C. et al. Transcriptomic classes of BCR-ABL1 lymphoblastic leukemia. Nat. Genet. 55, 1186–1197 (2023).

31. Dixon, J. R. et al. Integrative detection and analysis of structural variation in cancer genomes. Nat. Genet. (2018) doi:10.1038/s41588-018-0195-8.

32. Lee, S. H. R. et al. Prognostic and pharmacotypic heterogeneity of hyperdiploidy in childhood ALL. J. Clin. Oncol. 41, 5422–5432 (2023).

33. Haas, O. A. & Borkhardt, A. Hyperdiploidy: the longest known, most prevalent, and most enigmatic form of acute lymphoblastic leukemia in children. Leukemia 36, 2769–2783 (2022).

34. Rehn, J. A., O’Connor, M. J., White, D. L. & Yeung, D. T. DUX hunting—clinical features and diagnostic challenges associated with DUX4-rearranged leukaemia. Cancers (Basel) 12, 2815 (2020).

35. Tran, T. H. & Loh, M. L. Ph-like acute lymphoblastic leukemia. Hematology Am. Soc. Hematol. Educ. Program 2016, 561–566 (2016).

36. Boer, J. M. et al. BCR-ABL1-like cases in pediatric acute lymphoblastic leukemia: a comparison between DCOG/Erasmus MC and COG/St. Jude signatures. Haematologica 100, e354–e357 (2015).

37. Ofran, Y. & Izraeli, S. BCR-ABL (Ph)-like acute leukemia—Pathogenesis, diagnosis and therapeutic options. Blood Rev. 31, 11–16 (2017).

38. Liu, Z., Lee, D.-S., Liang, Y., Zheng, Y. & Dixon, J. R. Foxp3 orchestrates reorganization of chromatin architecture to establish regulatory T cell identity. Nat. Commun. 14, 6943 (2023).

39. Zylbersztejn, F. et al. The BMP pathway: A unique tool to decode the origin and progression of leukemia. Exp. Hematol. 61, 36–44 (2018).

40. Tsherniak, A. et al. Defining a cancer dependency map. Cell 170, 564-576.e16 (2017).

41. O’Byrne, S. et al. Discovery of a CD10-negative B-progenitor in human fetal life identifies unique ontogeny-related developmental programs. Blood 134, 1059–1071 (2019).

42. Corces, M. R. et al. Lineage-specific and single-cell chromatin accessibility charts human hematopoiesis and leukemia evolution. Nat. Genet. 48, 1193–1203 (2016).

43. Lee, D. D. & Seung, H. S. Learning the parts of objects by non-negative matrix factorization. Nature 401, 788–791 (1999).

44. Alexandrov, L. B. et al. Signatures of mutational processes in human cancer. Nature 500, 415–421 (2013).

45. Cho, Y.-J. et al. Integrative genomic analysis of medulloblastoma identifies a molecular subgroup that drives poor clinical outcome. J. Clin. Oncol. 29, 1424–1430 (2011).

46. Brunet, J. P., Tamayo, P., Golub, T. R. & Mesirov, J. P. Metagenes and molecular pattern discovery using matrix factorization. Proc Natl Acad Sci U S A 101, 4164–4169 (2004).

47. Meuleman, W. et al. Index and biological spectrum of human DNase I hypersensitive sites. Nature 584, 244–251 (2020).

48. Kotliar, D. et al. Identifying gene expression programs of cell-type identity and cellular activity with single-cell RNA-Seq. Elife 8, (2019).

49. Morgan, D. & Tergaonkar, V. Unraveling B cell trajectories at single cell resolution. Trends Immunol. 43, 210–229 (2022).

50. Tan, C. et al. NR4A nuclear receptors restrain B cell responses to antigen when second signals are absent or limiting. Nat. Immunol. 21, 1267–1279 (2020).

51. Prazma, C. M., Yazawa, N., Fujimoto, Y., Fujimoto, M. & Tedder, T. F. CD83 expression is a sensitive marker of activation required for B cell and CD4+ T cell longevity in vivo. J. Immunol. 179, 4550–4562 (2007).

52. Gebhardt, C., Németh, J., Angel, P. & Hess, J. S100A8 and S100A9 in inflammation and cancer. Biochem. Pharmacol. 72, 1622–1631 (2006).

53. Lin, Y. C. et al. A global network of transcription factors, involving E2A, EBF1 and Foxo1, that orchestrates B cell fate. Nat. Immunol. 11, 635–643 (2010).

54. Mansson, R. et al. Positive intergenic feedback circuitry, involving EBF1 and FOXO1, orchestrates B-cell fate. Proc. Natl. Acad. Sci. U. S. A. 109, 21028–21033 (2012).

55. Lin, H. & Grosschedl, R. Failure of B-cell differentiation in mice lacking the transcription factor EBF. Nature 376, 263–267 (1995).

56. Nechanitzky, R. et al. Transcription factor EBF1 is essential for the maintenance of B cell identity and prevention of alternative fates in committed cells. Nat. Immunol. 14, 867–875 (2013).

57. Kikuchi, H., Nakayama, M., Takami, Y., Kuribayashi, F. & Nakayama, T. EBF1 acts as a powerful repressor of Blimp-1 gene expression in immature B cells. Biochem. Biophys. Res. Commun. 422, 780–785 (2012).

58. Pongubala, J. M. R. et al. Transcription factor EBF restricts alternative lineage options and promotes B cell fate commitment independently of Pax5. Nat. Immunol. 9, 203–215 (2008).

59. Banerjee, A., Northrup, D., Boukarabila, H., Jacobsen, S. E. W. & Allman, D. Transcriptional repression of Gata3 is essential for early B cell commitment. Immunity 38, 930–942 (2013).

60. Roberts, K. G. et al. Targetable kinase-activating lesions in Ph-like acute lymphoblastic leukemia. N. Engl. J. Med. 371, 1005–1015 (2014).

61. Paulsson, K. & Johansson, B. High hyperdiploid childhood acute lymphoblastic leukemia. Genes Chromosomes Cancer 48, 637–660 (2009).

62. Enshaei, A., Vora, A., Harrison, C. J., Moppett, J. & Moorman, A. V. Defining low-risk high hyperdiploidy in patients with paediatric acute lymphoblastic leukaemia: a retrospective analysis of data from the UKALL97/99 and UKALL2003 clinical trials. Lancet Haematol. 8, e828–e839 (2021).

