## Supplemental Material for "Heterogeneity in chromatin structure drives core regulatory pathways in B-cell Acute Lymphoblastic Leukemia"

### Materials and methods.

#### Recruitment of pediatric B-ALL cohort and sample isolation.

Patients with leukemia were recruited at Rady Children's Hospital San Diego. Patients and/or their parents gave informed consent or assent for participation in this IRB-approved study (UC San Diego Human Research Protections Program #171672 and #130794) as appropriate for the patients' age. Peripheral blood or bone marrow samples were obtained from patients at the time of diagnosis or relapse.

Mononuclear cells were isolated from peripheral blood (PBMCs) or bone marrow biopsies (BMMCs). In brief, samples were diluted in a 1:1 volume ratio with Dulbecco's Phosphate Buffered Saline (DPBS, Gibco) with 2% Fetal Bovine Serum (FBS) and filtered through a 100µM cell strainer. Samples were then layered in a 1:2 volume ratio with 10mL of Lymphoprep (Stemcell Technologies) in a 15mL conical tube and centrifuged at 300g for 40 minutes at 4C in a swinging bucket centrifuge with the brake off. If more than 5mL of sample was available at this stage, this was split into multiple 15mL conical tubes. The mononuclear cells layer was then carefully pipetted off and washed twice with DPBS with 2% FBS at 300g for 5 minutes with the brake off. Cells were resuspended in PBS + 2%FBS and counted on an automated TC20 cell counter (Bio-rad) to measure the concentration of cells and estimate the fraction of viable cells. Cells were then aliquoted for different assays, with the goal of aliquoting 100,000 for each ATAC-seq experiment, 2-5 million for each Hi-C experiment, and 1-2 million for each RNA-seq experiment.

#### Hi-C Library Generation

For Hi-C assays, cells were resuspended in DPBS and fixed with 1% formaldehyde at room temperature for 10 minutes, and then quenched with 0.2M Glycine for 5 minutes. The cells were then pelleted and washed with PBS and the fixed pellets were flash frozen until ready for further processing. The Hi-C libraries were generated using the *in situ* Hi-C method[66] using the MboI restriction enzyme. Final library quality was initially checked by shallow sequencing on a MiSeq/Miniseq (Illumina), and if of sufficient quality was sequenced more deeply. For moderate depth Hi-C libraries, we sequenced using a NextSeq 500 machine (Illumina) with a goal of obtaining ~100 million final reads, while high depth samples were sequenced on a Novoseq 6000 (Illumina) with a goal of obtaining ~500 million reads.

#### ATAC sequencing library preparation.

ATAC-seq experiments for mononuclear cells isolated from the whole-blood or bone marrow of B-ALL patients were performed using the Omni-ATAC-seq method[67]. In each experiment, 1 x$10^5$ cells were centrifuged at 1000 x g for 10 min at 4 ºC. Following aspiration, a cell count of the supernatant was performed, the remaining cell number was calculated, and all further reagents in the protocol were titrated to this cell number.

Library construction was conducted using the Omni-ATAC-seq method. For every 5 x $10^5$ cells, nuclei were isolated in 50 ul cold ATAC-Resuspension Buffer (RSB) (10 mM Tris-HCl pH 7.4, 10 mM NaCl, 3 mM MgCl2) containing 0.1% NP40, 0.1% Tween-20, and 0.01% Digitonin, and pipet-mixed up-and-down at least 5 times. Nuclei isolation mix was incubated on ice for 3 exactly minutes, washed in 1 ml of cold ATAC-RSB containing 0.1% Tween-20 (but no NP40 or Digitonin) and centrifuged at 1000 x g for 10 min at 4 ºC. Nuclear DNA was tagmented in 50 ul Transposition mix (25ul 2 × TD buffer, 2.5ul transposase (100nM final), 0.5 ul 1% digitonin, 0.5 ul 10% Tween-20, 16.5 ul PBS and 5 ul diH2O), and incubated in a thermomixer at 37°C, 1000 x g for 30 min. Tagmented DNA was purified with Zymo DNA Clean and Concentrator-5 Kit (cat# D4014). Library amplification was performed using custom indexing Nextera primers from IDT in a 50 ul Kapa Hi Fi Hot Start PCR reaction (cat# KK2602). Following 3 initial cycles 1 ul of PCR reaction was diluted in 1 ml buffer (10mM Tris-HCl pH 8.0, 0.05% Tween) and 4 ul of each dilution were run in triplicate 20uL quantitative PCR (Kapa qPCR Library Quantitation Kit cat# KK4824) to calculate the optimum number of final amplification cycles. Library amplification was followed by SPRI size selection with Kapa Pure Beads (cat# KK8002) to retain only fragments between 80-1,200 bp. Library size was obtained on an Agilent Bio-Analyzer or TapeStation using a High Sensitivity DNA kit and factored into final Kapa qPCR results to calculate the final size-adjusted molarity of each library.

ATAC-seq libraries were pooled by size-adjusted molarity and sequenced to obtain a minimum of 30 to 50 million reads from each cell line in paired-end 42 x 42 configuration.

**RNA-seq library generation**

Purified patient samples from 1-2 million cells were pelleted and resuspended in Trizol (Invitrogen) and incubated for 5 minutes at room temperature. After incubation, samples were typically frozen at -80°C until ready for further processing. RNA was extracted using RNeasy Mini Kit (Invitrogen #74106) using Purelink on-column DNase I digestion (Invitrogen #12185010) to remove residual genomic DNA. RNA quality was evaluated using RNA tapestation (Agilent) and processed only if high quality RIN values were obtained. RNA libraries were generated using the TruSeq Stranded mRNA-sequencing kit (Illumina #20020594) according to the manufacturer's instructions. Libraries were sequenced using a paired-end 42bp configuration on a NextSeq 500 (Illumina) machine with a goal of obtaining ~25 million read-pairs per sample.

**Single Cell RNA-seq and RNA+ATAC-seq multiome experiments**

Single-cell RNA-seq and Single-cell RNA+ATAC-seq multiome datasets were generated from PBMCs isolated from patient samples upon sample collection. Single-cell RNA-seq was generated using the 10x genomics Chromatin Single Cell 3' Reagent Kit (v3.1 – Catalog PN-1000268). Single-cell RNA+ATAC-seq data was generated using the Chromatin Next GEM Single Cell Multiome ATAC+Gene Expression kit from 10x genomics according to the manufacturer's instructions (catalog # 1000283). Libraries were sequenced according to the manufacturer's instructions on the Illumina NextSeq 500 sequencer.

**RNA-seq alignment and processing.**

We aligned RNA-seq reads to the reference genome (hg38 or hg19) using STAR (2.5.3a) [68]. We filtered the aligned reads using mapping quality score cutoff of 20 and removed duplicated reads using samtools (1.9) [69]. We obtained read counts for gencode genes using featureCounts (1.6.3) [70] and for Fragments Per Kilobase per Million mapped read (FPKM) normalized the read counts using DESeq2 (1.30.0) [69]. We inferred cell type proportions for normalized RNA-seq data from B-ALL samples using xCell [71].

**Hi-C data alignment and processing**

Hi-C libraries were aligned with BWA-MEM[67] to the hg38 reference genome using a previously described in house pipeline[68]. PCR duplicate reads were removed using Picard MarkDuplicates. Processed Hi-C files (*.hic or *.cool) were generated using Juicer Tools[71] and Cooler[72], respectively.

Differential 3D chromatin contacts between transcriptional subtypes were identified using edgeR[73]. In brief, we extracted raw Hi-C interaction counts for all pairs of 25kb bins up to 2Mb in size. Each patient sample was considered as a replicate observation of a transcriptional subtype, excluding any subtypes with only 1 patient represented. Differences 25kb bin pairs were identified between transcriptional subtypes using an ANOVA-like test based on quasi-likelihood F-tests with FDR 5%.

Chromatin loops were identified using the Mustache[74] tool at 5kb resolution and filtered to retain loops with FDR <= 5%. Loops were merged across all patient samples, retaining only unique loops. For each loop, we used the distance normalization interaction frequencies for downstream analysis. The distance normalized values were quantile normalized across patient samples to account for potential technical biases on loop intensity. The final interaction frequency $F_{ij}$ of bin $O_{ij}$ was defined as the average interaction frequency of a box around each pixel ($F_{ij} = \frac{1}{9}\sum_{m=i-1}^{i+1}\sum_{n=j-1}^{j+1} O_{mn}$). Loops derived from adjacent pixels were further merged by averaging the $F_{ij}$ values for each adjacent pixel. After such processing, only loops with at least 1 sample showing a normalized observed/expected interaction frequency greater than 5 were retained for downstream analysis.

**Transcriptional subtype clustering and sample classification**

Gene expression data from our cohort were co-clustered with data from Gu et al.[68] to identify transcriptional subtypes using the approach described in Gu et al. with minor modifications. Specifically, a sample by gene read count matrix was quantile normalized using preprocessCore R-package. Batch effects were then removed using the combat tool. In examining the Gu et al. data, we observed batch effects related to both library preparation method (Stranded vs. non-stranded), read length, and data origin. Therefore, we considered eight total batches based on a mixture of these effects, with the eighth batch used for samples generated as part of this study. The eight batches specifically were "Salk", "Stranded_total_RNA_PE100bp", "Stranded_total_RNA_PE100bp_ECOG-ACRIN", "Stranded_total_RNA_PE75bp_P", "Stranded_total_RNA_PE75bp_S", "Unstranded_mRNA_PE100bp", "Unstranded_mRNA_PE50bp", and "Unstranded_mRNA_PE75bp". The top 1000 most variable genes across all samples were identified using Shannon Information Entropy content, and the normalized sample by gene matrix of these top 1000 most variable genes was the input for Uniform Manifold Approximation and Projection (UMAP) embedding and visualization.

To classify transcriptional subtypes of the samples generated as part of this study, we used the above identified top 1000 most variable genes and the samples from the Gu et al. study as training data for K-nearest Neighbor Classification (K=10). The sample labels for the training data were derived from the Gu et al. study. Expression of the top 1000 most variable genes in our dataset was then used as the test data to assign transcriptional subtype labels. We also tested using either the top 5 UMAP dimensions or all genes as training data, as well as varying the K parameter (10 or 30), and this had only very minor effects on classification.

**Identification of B-ALL selectively essential genes**

Identification of B-ALL selectively essential genes was performed using CRISPR/Cas9 knock-out screens from data downloaded from DepMap[40]. Essentiality scores represent the degree of depletion of gRNAs from the CRISPR/Cas9 knock-out screens for survival, with lower (more negative) essentiality scores representing more depleted guides and therefore more essential genes for survival in a given cell line. Essentiality scores were downloaded for all genes for all cell types in the DepMap database. The average essentiality for each gene was calculated among cell lines split into two groups, B-ALL cell lines and non-B-ALL cell lines. B-ALL cell lines were defined as the following: 697, ALLPO, BALL1, GRST, HAL01, HB1119, JM1, KASUMI2, KOPN8, LC41, MHHCALL2, MHHCALL3, MHHCALL4, MN60, MUTZ5, NALM16, NALM19, NALM6, P30OHK, RCHACV, REH, ROS50, RS411, SEM, SEMK2, SUPB15, SUPB8.

**Identification and analysis of structural variants from Hi-C data**

Hi-C data was used to detect structural variants using the hic-breakfinder tool as previously described[74]. Only structural variants that were called at the highest resolution (10kb) were kept for downstream analysis. Topologically associated domains were called as previously described on a merged Hi-C dataset merging data from all high-resolution patient samples. The number of SVs in each TAD was summed across samples. For gene expression changes near SV breakpoints, fold changes were computed for each gene in each sample by dividing each RPKM by the average RPKM across all samples. Windows at different sizes (500kb, 400kb, 300kb, 250kb, 200kb, 150kb, 100kb and 50kb) were made upstream from the SV breakpoint. Each gene fold change was classified as "affected" if the gene's TSS falls into the SV associated window in the same sample and classified as "unaffected" if not. The fraction of genes with fold-change greater than two were plotted at different window sizes.

**Correlation of variable chromatin interactions with ATAC-seq or RNA-seq data**

For identifying genes and open chromatin sites that correlate with variable chromatin interactions, we first intersect the anchor of each chromatin interaction with an ATAC-seq peak or gene transcription start site. As a Hi-C chromatin interaction has two anchors, this means that each Hi-C data value can be correlated with genes or peaks at either anchor. We focused on the 20,195 variable chromatin interactions for this analysis. We compared the Hi-C interaction frequencies at these sites with RPM normalized ATAC-seq data (for ATAC-seq analysis) or with RPKM normalized gene expression in the same patient. The transcription start site was defined as a window of -2000bp to +500bp from the annotated gene transcription start site. For permutation analysis, we shuffled the vector of ATAC-seq values or RNA-seq values, maintaining the observed data points but randomly assigning them to different patients to maintain the structure of the data. This permutation was performed 1000 times. To calculate an empiric False Discovery Rate, we first rank-ordered the correlations for all HiC-ATAC (or HiC-RNA) random correlations genome wide after each round of permutations. We then calculated the average correlation of rank. We then rank-ordered the observed correlations and calculated the number of observed correlations greater than or equal to that value and compared this with the number of random correlations greater than or equal to the observed value. The FDR is then the number of greater random correlations divided by the number of greater observed correlations.

**Analysis of ATAC-seq data from B-lineage development**

ATAC-seq data from B-lineage development was downloaded from Corces et al.[42] and O'Byrne et al.[41]. Reads were mapped to hg38 reference using BWA-MEM[74] and PCR duplicates removed with Picard. Peaks were called using the "hmmratac" function in macs3[75]. Peaks were merged across all individuals from all cell types into a single pan-B-lineage development set. ATAC-seq enrichment plots were produced using DeepTools[76]. Due to disparate numbers of peaks called per sample, we normalized the coverage for each sample using an RPM normalization based on the sum of the 10,000 strongest ATAC-seq peaks. To identify significantly different peaks in B-lineage development, we used edgeR[75] to identify differential peaks across lineages treating each individual as a replicate observation for each lineage. We considered peaks as differential if they showed a 4-fold increase in ATAC-seq signal relative to the mean of all lineages and if they had a corrected P-value of less than 0.01 (Benjamini-Hochberg).

**Identifying gene signatures using non-negative matrix factorization.**

We obtained gene expression data for B-ALL samples from Gu et al, 2019[76] and selected 1,266 out of 1,988 samples for analysis because these samples were sequenced using the same protocol (i.e., 100bp paired-end stranded total RNA-seq). We normalized the read counts using the variance stabilizing transformation from DESeq2 (1.30.0) [77] and calculated information content (i.e., Shannon entropy) for each gene using the entropy package[77] (1.2.1). Then, we selected the top 5,000 genes ranked based on their information content and analyzed the gene expression data using non-negative matrix factorization (NMF). We implemented the NMF algorithm from Brunet et al, 2004 using the NMF package (0.21.0) [75].

NMF decomposes the gene expression matrix into the product of two matrices: a basis matrix and a sample coefficient matrix. We tested factorization ranks (k) from 6 to 24 with random initialization. We computed cophenetic coefficients[78] and pairwise correlations between sample coefficients for each factorization rank. Based on these statistics, we used a factorization rank of K=11, which identified stable components (cophenetic coefficient ≥ 90%) while avoiding redundancy between components (no component pairs with Pearson correlation ≥ 0.2).

To derive the sample coefficient matrix for our cohort of B-ALL patients, we solved the equation $a*x = b$ using QR factorization. Here, a is the reference basis matrix, b is the log-normalized gene expression matrix for our samples for the same set of 5,000 genes, and x is the target sample coefficient matrix. All heatmaps were generated using the ComplexHeatmap Package (1.20.0).

**Gene Ontology Analysis**

We performed Gene Ontology enrichment analysis for Biological Processes (GOBP) using topGO[78] (2.34.0). To select non-redundant GOBP terms for each NMF signature, we grouped the top 25 GOBP terms (ranked by Fisher's Exact Test p-value) and grouped them by semantic similarity using GOSemSim[79] (2.8.0). We then selected the most significant GOBP term for each group.

**Single Cell RNA-seq and RNA+ATAC-seq multiome data analysis**

Single-cell RNA-seq and RNA+ATAC-seq multiome data were aligned and analyzed using Seurat[80]. For scRNA-seq, reads were aligned and Shared Nearest Neighbors were identified using the first 11 principal components (PCs). Clustering was perfomed using a resolution parameter of 0.5. Data was visualized using Uniform Manifold Approximation and Projection (UMAP) using the first 11 PCs as input. From the clustering, we distinguished cycling from non-cycling cell clusters using the MKI67 marker gene. We further excluded cells of non-B-cell origin using CD247 as a marker gene for T-cells/NK cells. Differential gene expression was called using “FindMarkers” comparing each B-ALL cluster against all other B-ALL clusters. Significant differential genes were called using FDR=5% and a fold change cut-off of 2-fold increased expression relative to other clusters.

For single-cell RNA+ATAC-seq multiome experiments, we analyzed the data with Seurat. The data was aligned and we used “FindMultiModalNeighbors” to generate a weighted shared nearest neighbor graph based on both RNA and ATAC-seq data. We specifically used Latent Semantic Indexing reduction using the 2:50 dimensions for ATAC-seq data reduction, and PCA using the first 50 PCs for RNA-seq. Clusters were identified using the SLM algorithm with the resolution parameter set to 0.25. Data was visualized using UMAP from the weighted nearest neighbor graph. We excluded cycling cells based on the marker gene MKI67 and macrophages by the marker gene FCER1G. Differential gene expression was identified using only the scRNA-seq aspect of the data based on the B-ALL clusters using “FindMarkers.” Differential genes were called if they had a fold-change of greater than 2 relative to other B-ALL clusters (FDR=5%).

**Alignment, quality control and analysis of ATAC sequencing data.**

We aligned ATAC-seq reads to the reference genome (hg38) using BWA-mem[81]. We filtered the aligned reads using mapping quality score ≥ 20 and removed mitochondrial and duplicate reads using Samtools(1.9) [75]. Then we identified ATAC-seq peaks using MACS2.0 with parameters as described in the ENCODE ATAC-seq pipeline (--shift -75 --extsize 150 --nomodel --call-summits --keep-dup all -p 0.01)[75]. Finally, we computed quality metrics i.e., transcription start site enrichment scores (TSSE scores) using the ChrAccR (0.9.11) package in R-4.0.1.

**Correlating chromatin accessibility with gene expression programs.**

To compare chromatin accessibility between samples, we divided the genome into 1kb bins and retained bins that overlapped with ATAC-seq peaks identified using MACS2.0 in at least one B-ALL sample. We then performed a Spearman Rank Correlation between ATAC-seq bins and Derived Coefficient for NMF components in R-4.0.1[66]. To understand biological processes impacted by the bins associated with NMF components, we analyzed them using the Genomic Regions Enrichment of Annotations Tool (GREAT) (https://great.stanford.edu).

**Motif enrichment analysis**

We compared the occurrence of human transcription factor binding motifs from the JASPAR2020 motif database correlated bins (i.e., foreground) to a random set of 10,000 high coverage ATAC-seq bins (i.e., background) for each cluster using findMotifsGenome.pl function in HOMER2. Motif enrichment bar plots were generated using ggplot2 (3.3.5) in R-4.0.1. A similar approach was used for identical motif enrichment at ATAC-seq peaks at loop anchors and across the body of loops, but the background regions were auto-generated by Homer to match the observed nucleotide content of the foreground ATAC-seq peaks.

**Survival Analysis of B-ALL samples from TARGET.**

To determine if NMF components are associated with survival, we analyzed gene expression and survival data for 204 B-ALL patients from the Therapeutically Applicable Research To Generate Effective Treatments (TARGET) program. We normalized the read counts using the variance stabilizing transformation from DESeq2 (1.30.0) and corrected the vst-matrix for batch effect (i.e. 2 batches dicentric or phase2) using *removeBatchEffect* function from R-package limma. Then, we derived the sample coefficient matrix for TARGET patients using QR factorization and assigned patients to NMF components as described previously. Finally, significance levels and hazard ratios for NMF components were calculated using the Cox Proportional Hazards model in R-4.0.1. Plots were generated using ggplot2 (3.3.5) in R-4.0.1.

**References**


63. Rao, S. S. P. *et al.* A 3D map of the human genome at kilobase resolution reveals principles of chromatin looping. *Cell* **159**, 1665–1680 (2014).
64. Corces, M. R. *et al.* An improved ATAC-seq protocol reduces background and enables interrogation of frozen tissues. *Nat. Methods* **14**, 959–962 (2017).
65. Dobin, A. *et al.* STAR: ultrafast universal RNA-seq aligner. *Bioinformatics* **29**, 15–21 (2013).
66. Li, H. *et al.* The Sequence Alignment/Map format and SAMtools. *Bioinformatics* **25**, 2078–2079 (2009).
67. Liao, Y., Smyth, G. K. & Shi, W. featureCounts: an efficient general purpose program for assigning sequence reads to genomic features. *Bioinformatics* **30**, 923–930 (2014).
68. Love, M. I., Huber, W. & Anders, S. Moderated estimation of fold change and dispersion for RNA-seq data with DESeq2. *Genome Biol* **15**, 550 (2014).
69. Aran, D., Hu, Z. & Butte, A. J. xCell: digitally portraying the tissue cellular heterogeneity landscape. *Genome Biol* **18**, 220 (2017).
70. Li, H. Aligning sequence reads, clone sequences and assembly contigs with BWA-MEM. *arXiv [q-bio.GN]* (2013).
71. Durand, N. C. *et al.* Juicer provides a one-click system for analyzing loop-resolution hi-C experiments. *Cell Syst.* **3**, 95–98 (2016).
72. Abdennur, N. & Mirny, L. A. Cooler: scalable storage for Hi-C data and other genomically labeled arrays. *Bioinformatics* **36**, 311–316 (2020).
73. Robinson, M. D., McCarthy, D. J. & Smyth, G. K. edgeR: a Bioconductor package for differential expression analysis of digital gene expression data. *Bioinformatics* **26**, 139–140 (2010).
74. Roayaei Ardakany, A., Gezer, H. T., Lonardi, S. & Ay, F. Mustache: multi-scale detection of chromatin loops from Hi-C and Micro-C maps using scale-space representation. *Genome Biol.* **21**, 256 (2020).
75. Zhang, Y. *et al.* Model-based analysis of ChIP-Seq (MACS). *Genome Biol* **9**, R137 (2008).

76. Ramírez, F. *et al.* deepTools2: a next generation web server for deep-sequencing data analysis. *Nucleic Acids Res.* **44**, W160–W165 (2016).

77. Hausser, J. & Strimmer, K. Entropy Inference and the James-Stein Estimator, with Application to Nonlinear Gene Association Networks. *J. Mach. Learn. Res.* **10**, 1469–1484 (2009).

78. Adrian Alexa, J. R. *TopGO*. (Bioconductor, 2017). doi:10.18129/B9.BIOC.TOPGO.

79. Yu, G. *et al.* GOSemSim: an R package for measuring semantic similarity among GO terms and gene products. *Bioinformatics* **26**, 976–978 (2010).

80. Hao, Y. *et al.* Integrated analysis of multimodal single-cell data. *Cell* **184**, 3573-3587.e29 (2021).

81. Houtgast, E. J., Sima, V. M., Bertels, K. & Al-Ars, Z. Hardware acceleration of BWA-MEM genomic short read mapping for longer read lengths. *Comput Biol Chem* **75**, 54–64 (2018).

**Supplementary Table.**

**Table S1:** Clinical characteristics of our B-ALL cohort.

**Table S2:** Hi-C sample sequencing and data quality

**Table S3:** Hi-C defined structural variants

**Table S4:** Fusion genes identified from RNA-seq

**Table S5:** Gene Ontology analysis of NMF associated genes

**Table S6:** ATAC-seq distal or intronic bins correlated with at-least one NMF gene signature

**Table S7:** Motif enrichment analysis for distal or intronic bins correlated with at-least one gene expression program.
